# Transposable Element Expression Predicts Prognosis in Acute Myeloid Leukemia

**DOI:** 10.1101/217299

**Authors:** Anthony Colombo, Timothy Triche, Giridharan Ramsingh

**Affiliations:** Keck School of Medicine of University of Southern California, Jane Anne Nohl Division of Division of Hematology and Center for the Study of Blood Diseases; Los Angeles, California, 90033.

**Author notes:** Corresponding authors and 1441 Eastlake Ave Los Angeles, CA 90033 USA.

## Abstract

Over half of the human genome is comprised of transposable elements (TE). TE have been implicated in cancer pathogenesis. Despite large-scale studies of the transcriptome in cancer, a comprehensive look at TE expression investigating its relationship to various mutations and its role in predicting prognosis has not been performed. We characterized TE expression in 178 adult acute myeloid leukemia (AML) patients using transcriptome data from The Cancer Genome Atlas (TCGA). We identified significant dysregulation of TE, with distinct patterns of TE expression correlated to specific mutations and distinct coding gene networks. TP53 mutated AML had a unique TE expression signature and was associated with significantly suppressed expression of TE and various classes of non-coding RNA. We identified 17 candidate prognostic TE transcripts that can classify AML subtypes as either high or low risk. These 17 TE were able to further sub-stratify low risk AML (based on mutational profile and coding gene expression) into favorable and unfavorable prognostic categories. The expression signature of the 17 TE was able to predict prognosis in an independent cohort of 284 pediatric AML patients, and was also able to predict time to relapse in an independent dataset of relapsed adult cases. This first comprehensive study of TE expression in AML demonstrates that TE expression can be used as a biomarker for predicting prognosis in AML and also provides novel insights into the biology of TP53 mutated AML. Studies characterizing its role in other cancers are warranted.

## INTRODUCTION

Approximately 50% of the genome is comprised of transposable elements (TE), including satellites, long interspersed nuclear elements (LINE), short interspersed nuclear elements (SINE), DNA/RNA transposons, long-terminal repeats (LTR) including endogenous retroviruses (ERV), and rRNA ^1^. Despite large-scale studies of genome and transcriptome over the last decade, the importance of TE in health and disease has been a focus of intense research only recently. TE have been implicated in the pathogenesis of cancer, but studies have mostly focused on their deleterious effects. For example, LINE1 ORF overexpression has been observed in many cancers suggesting increased transpositioning activity ^2–4^, which can lead to increased genomic plasticity and insertional mutagenesis. Recent analysis of TE expression in large cancer transcriptome database demonstrated increased activity of LINE1 in cancer ^5^. However, evidence for direct mutagenic effect of TE in cancer is sparse ^6^. Moreover, LTR, including ERV, are mostly defective in their transpositioning activity. Very recently, beneficial roles of TE have been described. Induction of their expression leads to the activation of the viral recognition pathway, enabling cancer cell death ^7,8^. In addition, TE have been shown to regulate coding gene function ^9–12^. Hence, they may indirectly alter the transcriptional networks to promote or inhibit cancer cell growth. This suggests that TE have complex and diverse functions in cancer, which has largely remained unexplored.

Prediction of prognosis using coding gene expression in cancer has been widely studied and this has resulted in development of many assays for clinical use. Hypomethylation of LINE1 element in the genome of cancer has been associated with prognosis in cancer ^13^, however the role of TE expression in predicting prognosis in cancer has not been explored comprehensively.

Regulation of TE expression remains poorly understood. Like coding genes, TE can be regulated both transcriptionally and post-transcriptionally ^14^. Epigenetic modifications, such as methylation of DNA and histones regulate TE expression, as do transcription factors such as ATRX, P53, and SIRT6 ^15–17^. However, the identification of transcriptional circuits and the cofactors involved in TE regulation remains incomplete.

The role of coding gene mutations in the alteration of gene network is well known, providing valuable information on the regulation of coding genes by genes mutated in cancer. However, how mutations affect transcription of TE has not been well characterized. By understanding the changes in TE expression with respect to specific mutations in cancer, we can gain novel insight into how TE expression is regulated by the genes that are mutated in cancer.

In this study, we performed a comprehensive analysis of the expression of TE in acute myeloid leukemia (AML) transcriptome. We analyzed the mutation specific alterations in expression of TE and characterized their expression pattern to transcriptional networks. We identified a TE expression signature that predicts prognosis in AML, paving way for the development of novel biomarkers for prognostication in AML.

## RESULTS

### Mutation specific dysregulation of TE expression in AML

Mutations in AML are associated with distinct alterations in the expression of coding genes ^18^, which provides insight into gene regulation. In order to investigate the effect of mutations on the expression of TE and identify possible mechanism for regulation of TE in the context of AML, we investigated the relationship between specific mutations and expression of TE in AML. We analyzed the transcriptome from 178 AML patients in the cancer genome atlas (TCGA) using *Arkas* ^19^, an RNA sequence analysis pipeline that provides detailed annotation information for TE and ENSEMBL non-TE (non-TE) transcripts. The non-TE includes protein-coding genes, pseudogenes, long non-coding RNA (lncRNA), and short non-coding RNA.

The TE and non-TE were normalized together using voom^15^ log_2_ counts per million (CPM) expression. We used the multivariate empirical Bayesian linear model to study the effect of various mutations on the expression of TE ^18^. For this, we used the mutational status of 178 AML patients as independent predictors and TE expression as response. Hierarchical filtering, coupled with Benjamini-Hochberg (BH) false discovery rate (FDR) q.value threshold of 0.05, identified the total number of significantly (BH q.value ≤0.05) altered expression of TE transcripts (AE-TE) with respect to each mutation. TP53 mutation was associated with the most number (49) of AE-TE, with 40 showing down regulation (Figure 1A). This was followed by NPM1 mutation (36), with 20 showing down regulation. Mutations in Cohesin complex (SMC1A, STAG2, SMC3) were associated with the 3^rd^ most number of AE-TE, with SMC1A mutation being associated with 28 AE-TE (mostly exhibiting down regulation). RUNX1, inv (16) and MLL gene rearrangement were mostly associated with up regulation of TE. Although methylation has been shown to regulate the expression of TE ^7,8,17^, DNMT3A and TET2 mutations instead showed the lowest number of AE-TE.

**Figure 1:**
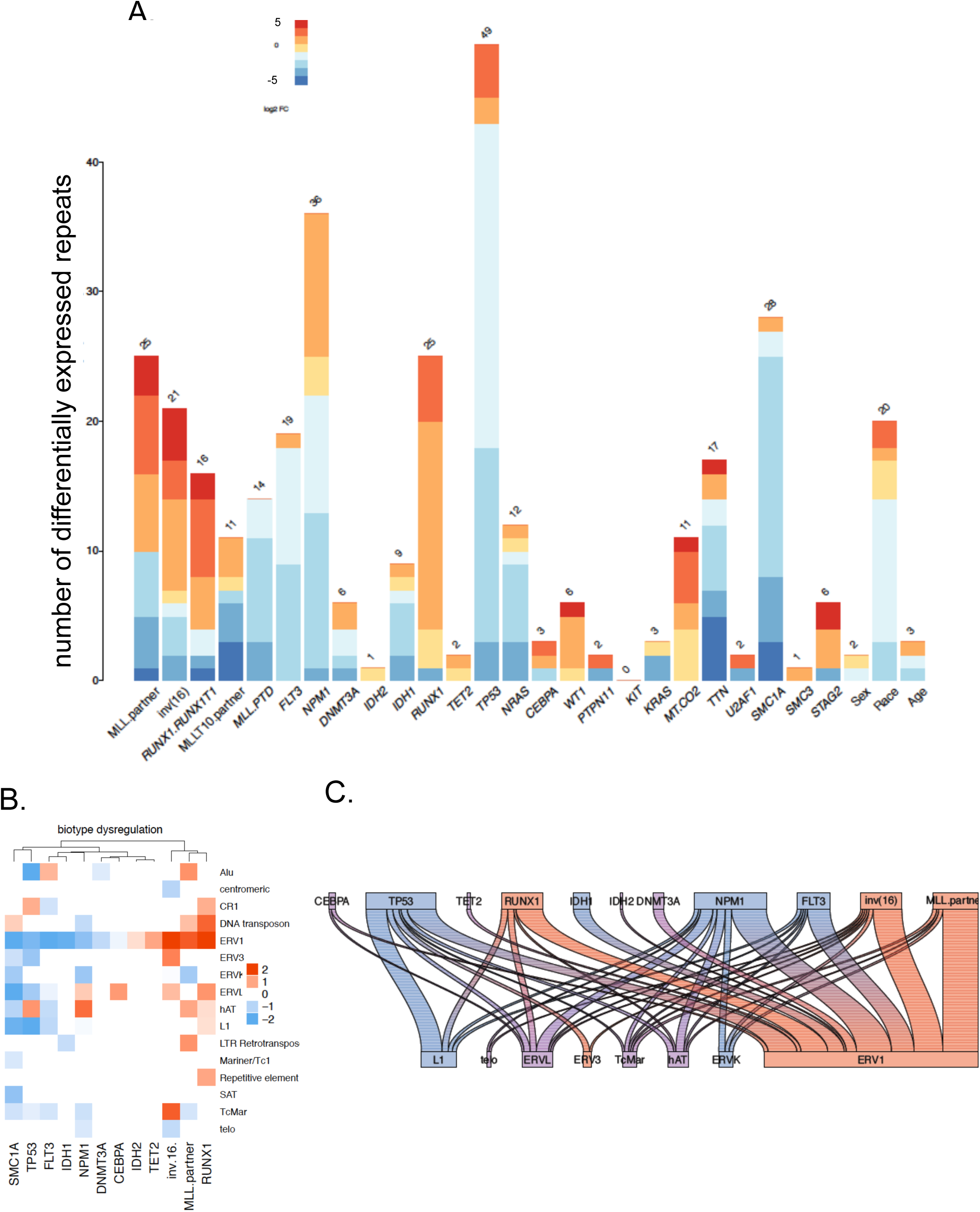
Mutation specific alteration in expression of transposable elements. **A)** *The number of TE altered in expression (AE-TE) with respect to specific mutation.* (Hierarchical test FDR <0.05; BH adjusted; n=178). **B)** TE biotype dysregulation with respect to specific mutation. Statistically significant coefficients total TE biotype sum identified from multiple regression per mutation (hierarchical test FDR <0.05; BH adjusted; n=178). **C)** *A summary river-plot of mutation (top axis) specific dysregulation of TE biotypes (sum of significant coefficients corresponding to TE) (bottom axis).* Red is estimated up-regulated, and blue is estimated down-regulated; purple indicates an even mixture of both up and down regulated transcripts corresponding to the respective TE biotype (hierarchical test FDR <0.05; BH adjusted; n=178).

TE biotypes exhibited specific alteration patterns with respect to mutational status (Figure 1B and 1C). LINE1 and ERVK were mostly down regulated with respect to mutations, whereas ERV3 and ERV1 exhibited up-regulation. ERVL showed both up and down regulation with respect to mutational status. Further, specific TE biotypes such as ERV1, ERVL, and L1 predominantly exhibited down-regulation with respect to TP53 mutation, and up-regulation with respect to MLL. partner rearrangement. These findings demonstrate a non-uniform regulation pattern of the expression of various TE transcripts and biotype predicted by mutation status, suggesting that that the common mutations in AML are associated with a distinct pattern of alteration in TE expression.

### Correlating the transcript network with the expression of TE

Regulation of TE expression and its downstream effects are not fully understood. TE are key regulators of coding gene expression ^1,9,10,12,20^. They are known to activate the interferon pathway, induce DNA breaks and DNA damage response. Distinct TE transcripts possibly mediate these varied effects. In order to gain insight into this, we correlated the normalized expression of TE biotypes with transcript networks. For this, the similarly expressed non-TE transcripts were grouped together, forming transcript modules ^21^ (Y-axis in Figure 2). These modules were then correlated with the expression of specific TE biotypes (X-axis in Figure 2). The number of non-TE transcripts that were altered in expression within each module, predicted by various mutations was measured. Mutations with high total number of significant altered expression of non-TE for each module were depicted (left column of Y-axis). This 3-way correlation matrix provided detailed information on the association between mutations, transcript networks, and the expression of various TE biotypes in AML.

**Figure 2:**
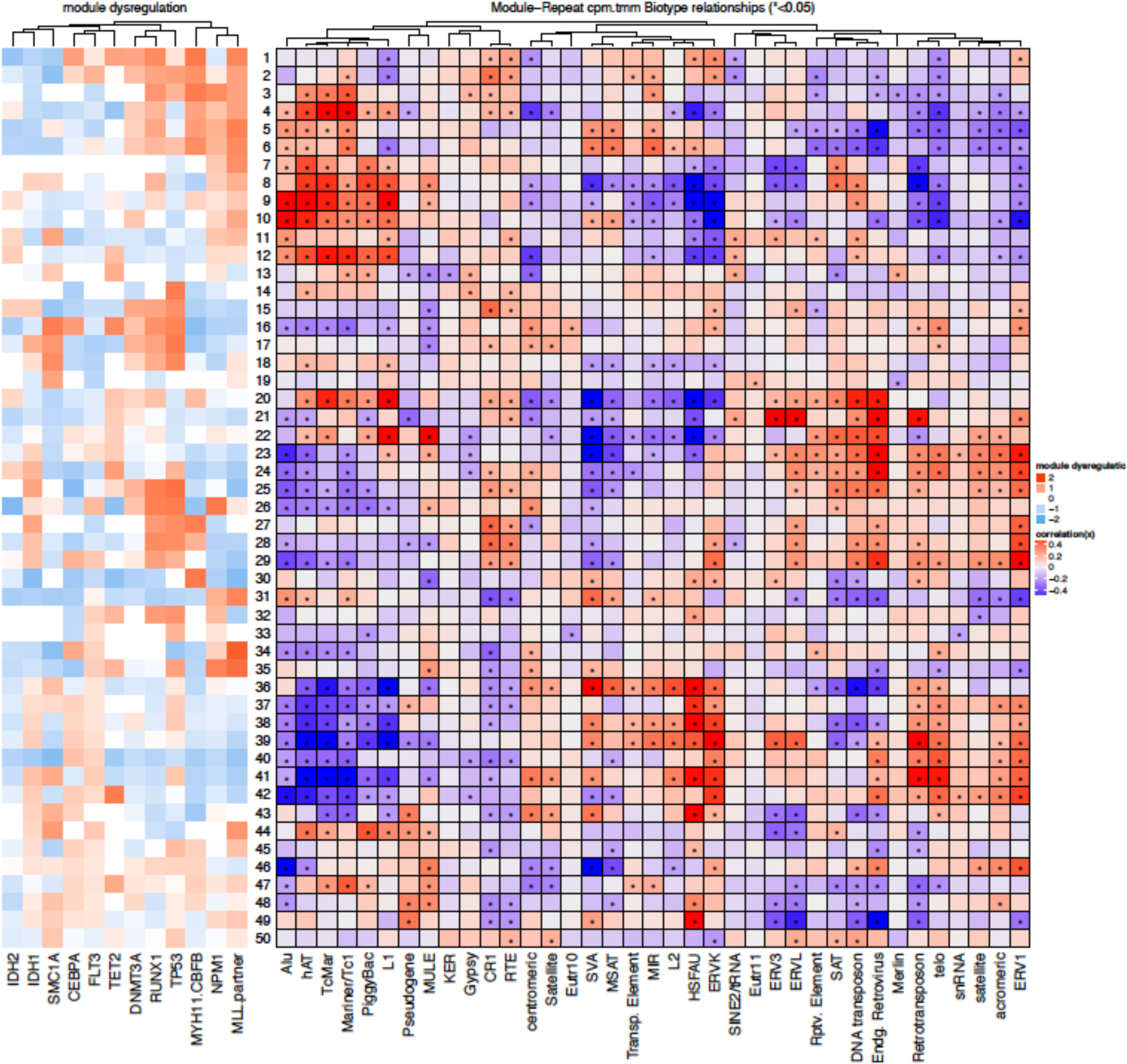
Correlating the transcript network with the expression of TE. Y-axis represents transcript 'modules' constructed by identifying non-TE transcripts based on the co-expression patterns. The X-axis denotes canonical TE biotypes used for correlating them. The centre figure represents the correlation matrix for the normalized gene 'module' expression and the TE type. * indicates significant associations (pearson correlation p.value ≤ 0.05). The panel to the left of the Y-axis depicts significant (hierarchical test FDR <0.05; BH adjusted; n=178) average dysregulation of the non-TE transcripts corresponding to each network module.

We observed that the TE biotypes formed distinct clusters based on its association with non-TE transcript modules, indicating diversity among TE biotypes. L1 was clustered furthest from ERV biotypes. Diversity was also observed among various ERV biotypes. For instance, the distinct sub-clusters that contained ERV1 held large distance from ERV3, ERVL and ERVK, indicating distinct association patterns. Though these findings do not prove direct functional relationship between TE biotypes and the transcript networks, they support diverse functionality of each TE subtype.

### Transcript network analysis in TP53 mutated AML exhibits unique signature

TP53 was associated with the most alteration in the expression of non-TE transcripts in the most number of transcript modules (Figure 2). Interestingly, TP53 was also associated with the highest number of alterations in expression of TE transcripts and non-TE (Figure 1A and Supplement Figure 1). In order to gain further insight into the association between non-TE network and TE in TP53 mutated AML, we generated a TP53 mutation specific network analysis (Figure 3A) using significantly altered non-TE (AE-non-TE) (Supplement Figure 1A) and AE-TE (Figure 1A) with respect to TP53 mutation. TP53 mutation was associated with 6809 AE-non-TE and 40 AE-TE (BH q.value ≤0.05, minimum absolute log-fold-change 0.2). Of the 6809 TP53 mutation specific AE-non-TE, 3918 were clustered into 7 modules with sufficient connectivity metrics as previously described ^21^. The 40 AE-TE were correlated to the principle component (PC) 1 of each 7 modules in the TP53 specific-network. Interestingly, the module 1 of the TP53 mutation specific network predominantly non-protein-coding transcripts: (722 (58%) lncRNAs, 157 pseudogenes (13%), 45 short non-coding RNAs (sncRNAs) (4%), and only 320 protein coding (26%)) (supplementary Table 3). It was also the only down regulated module with respect to TP53 mutation. In comparison, the other 6 modules had far less non-protein-coding RNA transcripts: modules 2, 3, 4, 5, and 6 held more than 100 transcripts in their modules and contained 703 (82.5%), 635 (74.6%), 377 (95%), 377 (99%) and 134 (95.7%) protein-coding genes respectively. Module 1 was also the only module to be statistically correlated to the TP53 specific AE-TE expression: 37/40 AE-TE had a statistically significant positive correlation to the expression of module 1. The non-coding RNAs in module 1 (lncRNAs, pseudogenes, and sncRNAs) were down-regulated with respect to TP53 mutation and were significantly positively associated with the expression of TP53 specific AE-TE. These two independent analyses associating the expression of TE and non-TE, in the context of TP53 mutation, showed a correlative link between different non-coding RNA classes with a common pattern of dysregulation in TP53 mutated AML. This may indicate that the mechanism of dysregulation of lncRNA, sncRNA and psedogenes may have a common link to the dysregulation of TE expression in the context of TP53 mutation in AML.

**Figure 3:**
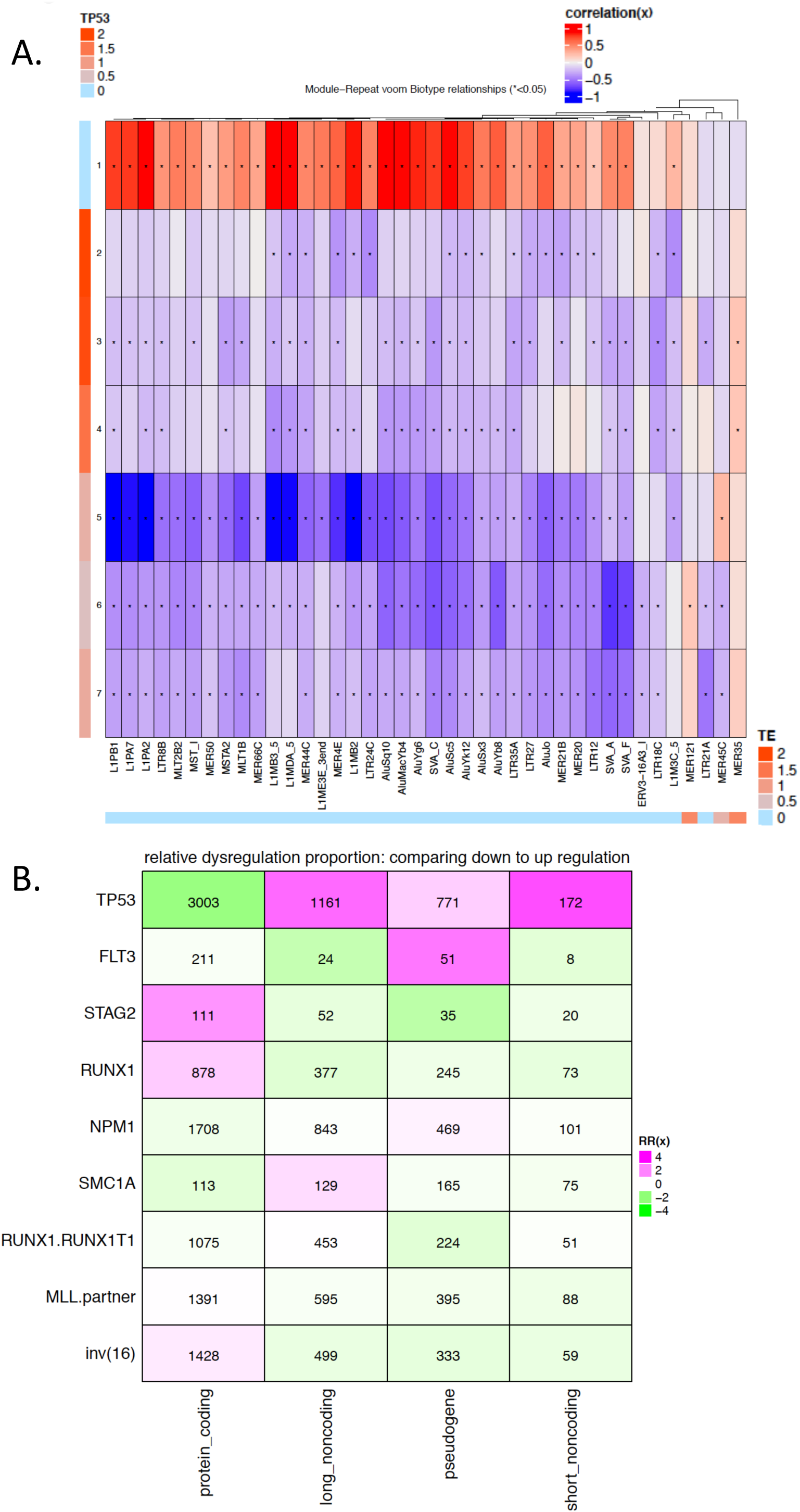
Transcript network analysis in TP53 mutated AML. **A)** *TP53 specific sub-network construction using TP53 specific dysregulated non-TE transcripts (AE-non-TE) to TP53 specific AE-TE.* The modules were constructed using the AE-non-TE as in Figure 2. To the left of Y-axis we represent the estimated altered expression of non-TE corresponding to each module (adjusted p.value <0.05; Benjamini-Hochberg adjustment method, minimum logFC 0.2). The x-axis depicts AE-TE predicted by TP53 dysregulation (adjusted p.value <0.05; Benjamini-Hochberg adjustment method, minimum logFC 0.2). The bottom x-axis annotation bar depicts the estimated altered expression per TE. The cells in the association table indicate the correlation of expression between the TE biotype and the PC1 of the module (pearson correlation statistical significant (“*”) p.value<0.05). **B)** *Comparison of dysregulation of non-TE classes between different mutations.* Mutational dysregulation relative proportions per non-TE class. The y-axis depicts each mutation used to measure significant dysregulation of non-TE (adjusted p.value <0.05; Benjamini-Hochberg adjustment method, minimum logFC 0.5). The x-axis depicts each class. The number in each cell are the total number of significant altered expressed of the non-TE transcripts in each class corresponding to each mutation. Purple indicates an increase in the proportion of suppressed non-TE transcripts for each class. Green indicates an increase in the proportion of activated non-TE transcripts similarly.

We sought to understand if this pattern of dysregulation of non-coding RNA classes is unique to TP53 mutation. For this, we tested the rates of dysregulation of different classes of non-TE in other mutations and compared it to TP53 mutation (Figure 3B). We measured the ratio of transcripts predicted to be down-regulated to that of up-regulated within each class of non-TE (protein coding, lncRNA, pseudogenes and small-ncRNA) for each mutation. TP53 mutation, which was associated with the mot dysregulation of both TE and non-TE transcripts, exhibited high level of suppression of non-coding non-TE classes and activation of protein coding-RNA. Other mutations were primarily associated with up-regulation of non-TE non-coding RNA classes.

These findings indicate that TP53 mutation in AML was associated with a unique signature of suppression of various classes of non-protein coding RNA and that this was linked with suppression of TE.

### Prediction of prognosis using TE expression in AML

Expression of coding genes and non-coding genes such as microRNAs has been shown to predict the survival of many cancers including AML ^22,23^. We investigated whether expression of TE can similarly predict prognosis in AML and if so, which TE are associated with good and bad prognosis. For this, the transcriptome from the 178 adult AML patients in TCGA was split into training and a test cohort. Initial discovery of the TE associated with prognosis was done in the training cohort and subsequently validated in the test cohort. In addition, further validation was performed in an independent 284 pediatric AML ^24^, and 19 relapsed adult AML ^25^ patients.

We randomly selected 99 (55%) patients into a training cohort, which ensured proportions of gender and racial factors were well balanced in each split of the full data. We performed a forward selection of TE transcripts that predict survival using univariate Cox regression. The forward selection used an unpenalized univariate Cox proportional hazard model with a significance threshold of 0.025 and 10-fold cross validation ^18^. This identified 17 TE associated with prognosis (TEP, Figure 4A). Following the identification of the TEP from the training cohort, we tested its ability to predict prognosis using the test cohort (79, 45% of patients). A multivariate Cox proportional hazard model using the TEP showed that the 17 TE covariates statistically distinguished patients with good and poor prognosis (Figure 4B, log-rank test p.value = 0.0041, score log-rank test=0.0027, wald test=0.0035). Upon testing the proportional hazards of the Cox model assumption using two-sided p-values, the 17 TEP did not violate the proportionality assumption. A 3-fold cross validation yielded a mean correspondence index (c-index) ^26–28^ of 0.55. The influence of gender and race was not associated to the TEP risk classification groups (gender Chi-square p.value=0.41; race Chi-square p.value=0.99).

**Figure 4:**
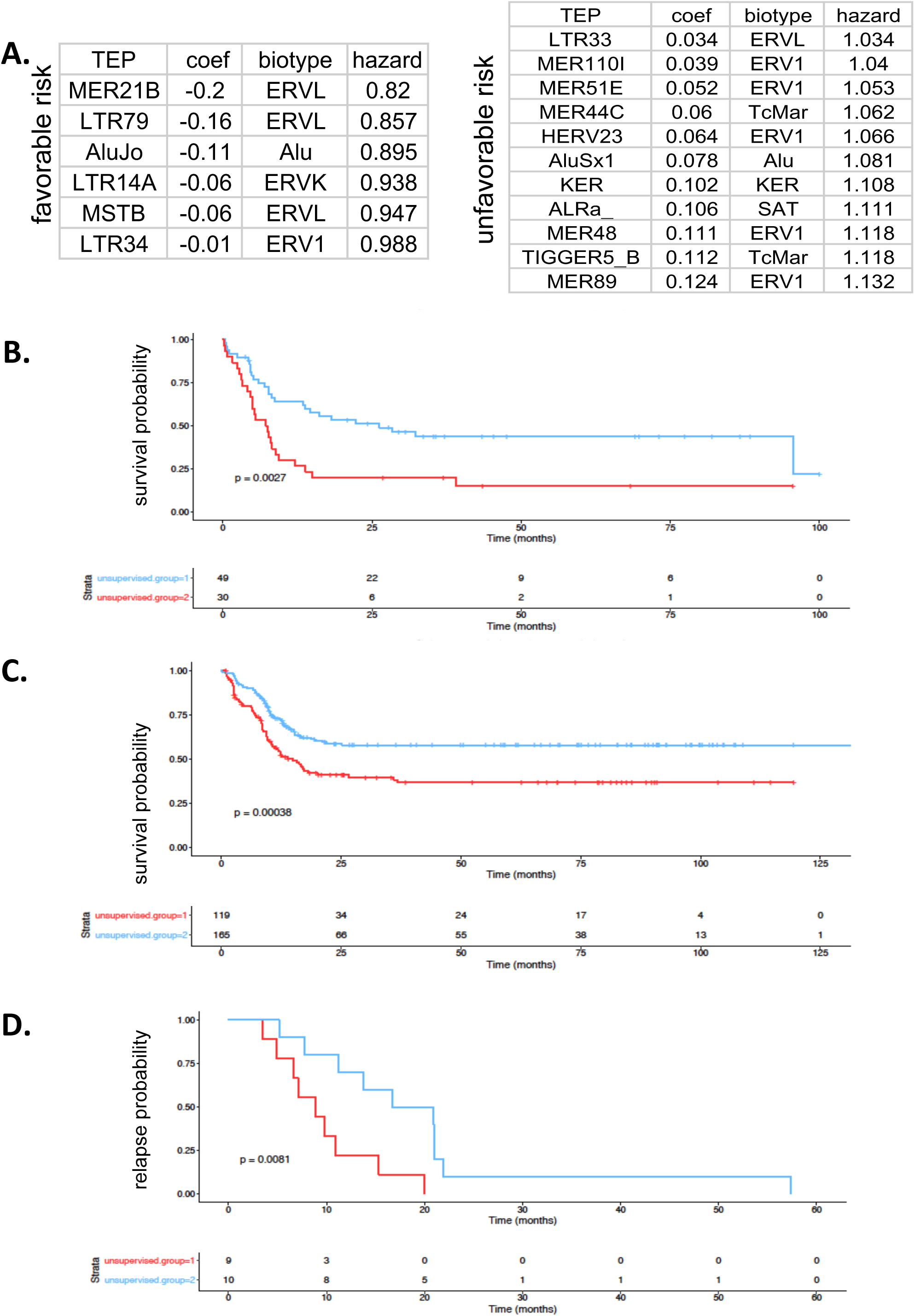
Prediction of prognosis using TE expression in AML. **A)** *TE transcripts identified to predict prognosis in AML (TEP).* The left table shows favorable risk TEP, estimated coefficients, biotype, and hazard estimates used in validation. The right table similarly shows unfavorable risk TEP. **B)** Validation of TEP in TCGA in the test cohort (N=79). Blue is favorable risk, and red is unfavorable risk. The y-axis is survival probability and the x-axis is time in months of 2 unsupervised patient risk classification groups determined by prognostic summary values (log-rank-test, score log-rank-test, and Wald test p.value <0.05). **C)** *Validation of the TEP in pediatric AML using TARGET data set (N=284).* Unsupervised classification groups were determined (log-rank-test, score log-rank-test, and Wald test p.value <0.05, n= 284) using the TEP. **D)** *Validation of the TEP in adult relapsed (N=19)*. The y-axis represents relapse probability, and the x-axis time in months. Unsupervised risk classification was similarly determined (log-rank-test, score log-rank-test, and Wald test p.value <0.05) using TEP.

In order to further confirm the validity of the TEP in predicting prognosis in AML, we used 2 independent cohorts (284 pediatric AML from TARGET and 19 relapsed adult AML). The RNA-seq data from these independent cohorts was processed using *Arkas* as previously described. Using the method described above to generate compound covariate summary values for risk estimates using the TEP expression and the corresponding Cox hazard estimates, we were able to stratify the 284 pediatric AML patients ^24^ into good and poor risk (Figure 4C, log rank test p.value=4.95e-04, score log rank test p.value = 3.79e-04, Wald test p.value=4.66e-04, N=284). Of the 17 TEP identified in TCGA, 16 were expressed in TARGET; AluSx1 was not sufficiently expressed in TARGET with at least 2 read counts across 284 samples. The log-rank test on the multivariate Cox model using the 16 TEP, without compounding hazard estimates into prognostic patient scores, revealed a statistically significant difference from a null model (log-rank test p-value=3.21e-05). Testing the Cox proportional hazards assumption using two-sided p-values showed all 16 TEP alpha levels greater than 0.05. After 5-fold cross validation, the c-index average across all folds was 0.584. This confirmed the validity of the 16 TEP in pediatric AML.

A second independent validation was similarly performed on a cohort of relapsed AML ^25^. Of the 17 TEP identified in TCGA, 14 TEP were expressed in the relapsed cohort; LTR33, KER, AluSx1 were not expressed higher than 0.2 across 19 samples. The TEP were able to stratify patients based on the differences in time to relapse between risk groups (Figure 4D, log rank test p.value=7.17e-04, score log rank test p.value =2.26e-04, Wald test p.value = 2.76e-03, N=19). The patient risk groups were formed using unsupervised clustering of prognostic summary values determined by compounding hazard estimates and corresponding TEP expression values. Upon examination, the Cox proportionality assumption was not violated using two-sided p-values. After 3-fold cross validation yielded a c-index of 0.57. These results indicated the robustness of the discovery algorithm that predicted prognosis using TE expression in a large cancer cohort.

We then wanted to identify whether the TEP would independently improve the mutation based risk stratification, which is a conventionally used biomarker for prognostication in AML. For this, we first used a Cox regression analysis and stratified the good-risk (N=99) and poor-risk (N=79) AML cohort using only the mutational signature. The TEP were then used to sub-stratify the low-risk and poor-risk groups identified by mutational status. In the mutation based low-risk group TEP signature reclassified 39/99 patients to a higher-risk group (Figure 5A, log-rank test p.value=0.0233, score log rank test p.value =0.0203, Wald test p.value=0.0222, N=99, Cox Assumption was not violated using two-sided p-values). After 3-fold cross validation, the c-index score of the TEP was on average 0.611 over all folds. Similarly the TEP expression signature was able to independently sub-stratify mutationally ‘good-risk cohort’ into better (40/79) and worse (39/79) groups (Figure 5B, log-rank test p.value=1.07e-03, scale log-rank test p.value=8.33e-04, wald test p.value=1.11e-03, Cox proportionality was not violated using two-sided p.value, 3 fold-CV c-index score=0.503).

**Figure 5:**
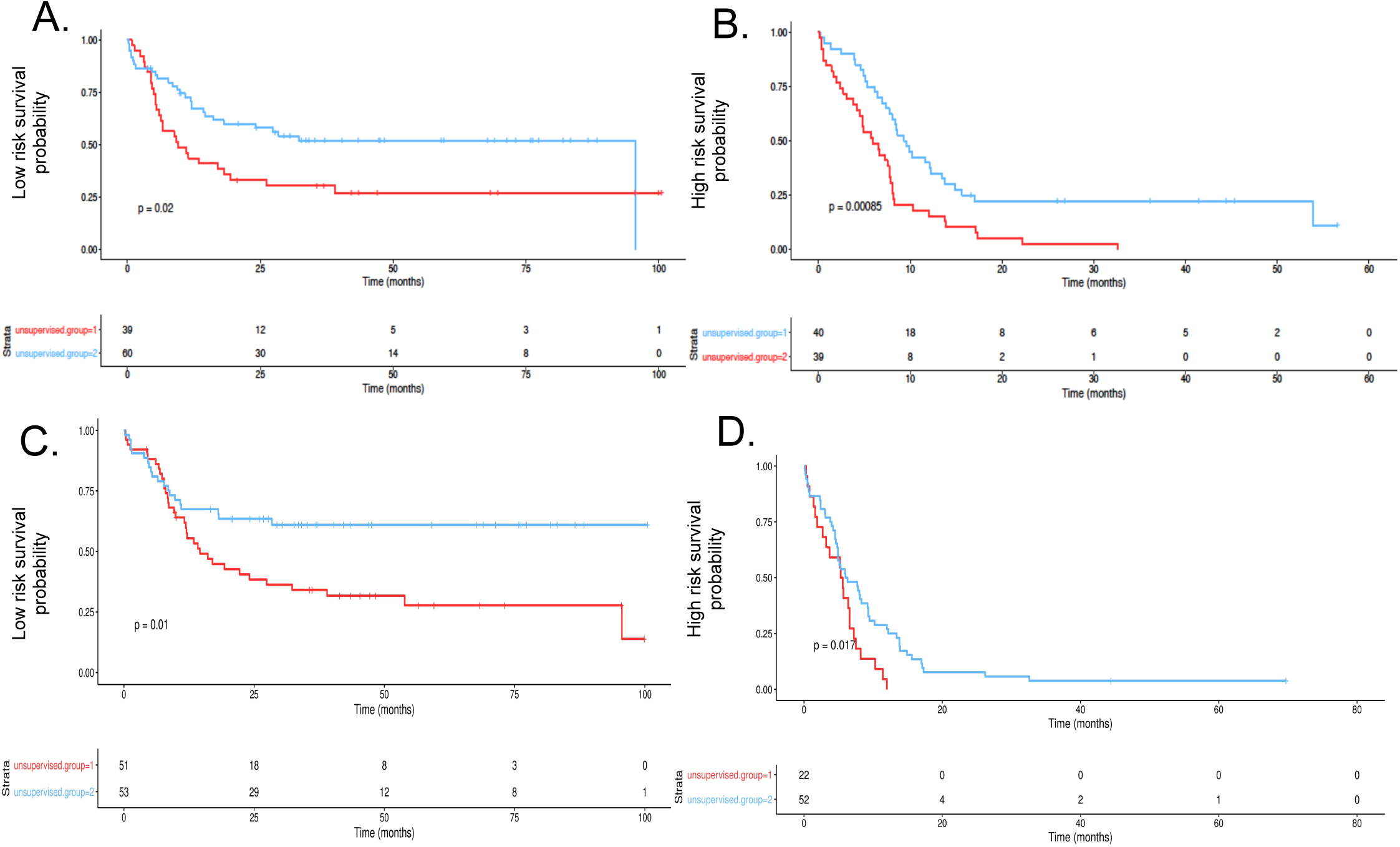
Utility of TE in improving mutation based and coding gene expression based risk prediction. **A)** *Improvement in prognostication of mutation based low-risk group (N=99).* Sub-stratification using 17 TEP (log-rank-test, score log-rank-test, and Wald test p.value <0.025, n= 99) of this cohort is represented. Favorable risk is blue, and unfavorable risk identified in red. B) *Improvement in prognostication of mutation based high-risk group (N=79).* Similar sub-stratification using 17 TEP (log-rank-test, score log-rank-test, and Wald test p.value <0.0125). C) *Improvement in prognostication of coding gene expression based low-risk group (N=83).* Low risk patients initially identified by using expression of 409 non-TE prognosticators were sub-stratified using TEP (log-rank-test, score log-rank-test, and Wald test p.value <0.025). **D)** *Improvement in prognostication of coding gene expression based high-risk group (N=95).* High-risk patients initially identified by using expression of 409 non-TE prognosticators were sub-stratified using TEP (log-rank-test, score log-rank-test, and Wald test p.value <0.0125, n= 95).

Similarly, TEP were able to independently sub-stratify risk classifications based on coding gene expression. The 17 TEP identified 41 poor risk patients from a pool of 83 good-risk patients identified based on gene expression (Figure 5C, log-rank test p.value=0.0173, scale log-rank test p.value=0.0163, wald test p.value=0.0195, Cox proportionality was not violated using two-sided p.value, 3 fold-CV c-index score=0.495). And, the 17 TEP identified 47 better risk patients from a pool of 93 poor-risk patients identified by gene expression Figure 5D, (log-rank test p.value=2.61e-03, scale log-rank test p.value=2.17e-03, wald test p.value=2.54e-04, Cox proportionality was not violated using two-sided p.value, 3 fold-CV c-index score=0.457).

Overall, this indicated that the TEP could provide robust independent prognostic value and can improve the prognostic ability obtained by either mutational status or coding gene expression in AML.

### Risk-stratification of AML based on a combination of mutations and TEP expression

In the previous analysis we showed that both mutations and TEP were able to risk-stratify AML patients and that TEP added independent prognostic value to mutation based risk stratification. We hence developed a composite model combining the prognostic value of both the mutations and TEP expression to understand the relative effect of mutations and TEP in various mutational sub-categories of AML. Using mutations alone the 178 AML patients were classified in to 99 low-risk and 79 high-risk patients. This risk stratification based placed most of DNMT3A, TP53, RUNX and FLT3 mutations into high-risk category and NPM1 mutation and inv (16) into low risk category.

A PC analysis, using the expression of TEP as vectors, was done on the 178 patients, identifying patients with and without specific mutations (Figure 6A). This provided information on the prognostic weight of the mutations relative to the effect of TEP. The results indicated that certain mutations (TP53, NPM1) showed significant deviation of the center to the high-risk 3-dimensional space, suggesting that these mutations by themselves provided strong prognostic value. DNMT3A, FLT3 mutations and inv (16) did not show significant deviation from the center, suggesting that patients with those mutations comprised of prognositically diverse group. We predicted that in these patients TEP would provide significant additional prognostic information.

**Figure 6:**
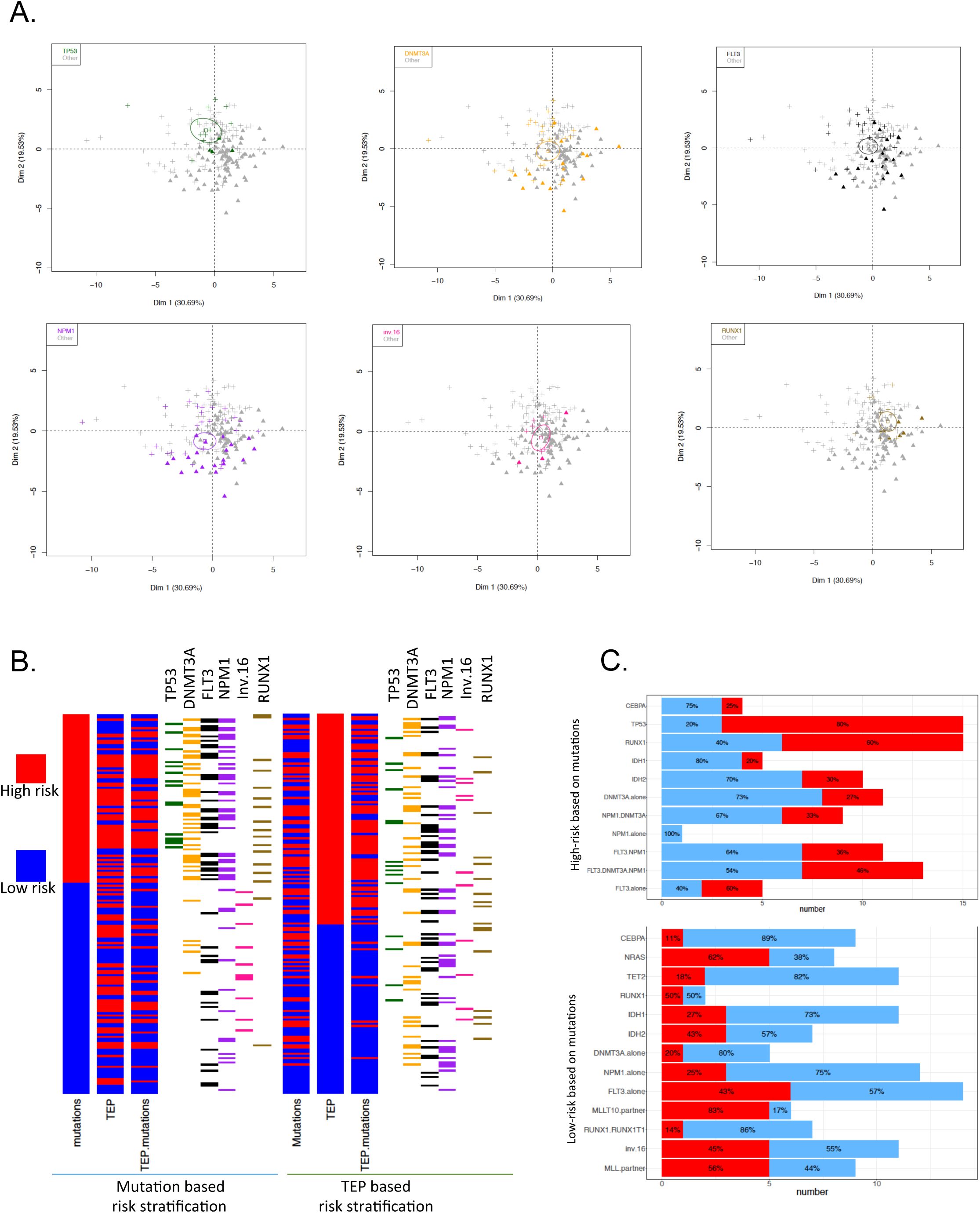
Risk-stratification of AML using Mutational profile and TEP. **A)** *Principle component analysis of 178 AML patients from TCGA using TEP expression.* In each panel, patients were classified based on presence or absence of the mutation and identified the location of the mutated patients in the 3-dimensional space. The solid triangles (‘+’) represent TEP risk categorization of low risk whereas crosses (‘+’) indicate high risk. Patients with mutations are represented in the respective color. **B)** *AML patients classified based on mutational profile vs. TEP.* The mutation based risk classification is depicted on the left. Each bar plot (3) depicts independent risk classification models using covariates such as mutations, 17 TEP, or mutations plus 17 TEP. High risk (red) and low risk (blue) represents patient’s risk classification. The TEP based risk stratification (right-most) similarly depicts bar plots (3) for each separate model. **C)** *Risk-adjustment based on TEP among different mutations in AML.* The top panel represents the mutational profile based high risk reclassified using TEP. TEP sub-stratified 79 patients, red indicates both models preserved high risk classification, blue indicates TEP sub-stratified the mutational classification to the TEP adjusted low-risk. In the bottom, we similarly depict the mutation profile of the 99 mutation based low-risk cohort sub-stratified by TEP. Blue indicates that both classification models preserved the patient as low-risk, and red indicates the TEP updated the mutational based low-risk to high-risk. Each bar plot depicts the percentage in each preserved/updated risk-stratification model.

TEP expression re-classified 39/99 mutation based low-risk and 40/79 mutation based high-risk patients to worse and better prognosis respectively (Figure 6B). We analyzed the mutational categories of patients that were reclassified in these groups (Figure 6C). In the mutation based high-risk group, a large majority of patients with DNMT3A mutations, DNMT3A plus NPM1 mutations, CEBPA mutations, FLT3 plus NPM1 mutations (73% of patients with DNMT3A mutation, 67% of DNMT3A plus NPM1 mutation, 75% of CEBPA mutation, and 64% of FLT3 plus NPM1 mutations) were re-classified as low-risk. In contrast, the TEP did not have a huge impact on reclassifying TP53 mutated patients identified as high-risk based on mutation profile (only 20% re-classified as low-risk). The low-risk patients (identified using mutational profile) were similarly reclassified to high-risk patients using TEP. Here, we observed that TEP significantly influenced re-classification to a higher-risk group in patients with mutations in NRAS, FLT3, MLL rearrangement, and inv (16).

This suggested that AML patients with DNMT3A, CEBPA or FLT3 mutations, MLL gene rearrangement and inv (16) constitute a diverse group with regards to their prognosis and that TEP provides significant prognostic information in these common subtypes of AML. The utility of TEP in re-classifying TP53 mutated AML and NPM1 mutated AML was minimal, suggesting that TP53 mutation and NPM1 mutation conferred strong risk prediction irrespective of the TEP expression status. These findings demonstrate that by incorporating the expression of 17 the TEP with the mutational status, we can significantly improve the ability to predict prognosis in AML, particularly in patients with common mutations such as DNMT3A, FLT3 mutations, MLL reaarnagement and inv (16).

## Discussion

This is the first study to comprehensively characterize the expression of TE expression at the transcript level, demonstrating novel roles for TE in cancer. Our study shows a strong association between TP53 mutation and suppression of TE in AML, a novel finding. Expression of TE is known to activate the viral recognition pathway ^7,8^. Though suppression of TP53 has been previously shown to activate the expression of TE and induce suicidal interferon response ^17^, our study showed that TP53 mutated AML exhibited features of TE suppression. Interestingly, we also observed suppression of several other classes of non-coding RNAs, including lncRNA and pseudogenes, many of which form dsRNA, in TP53 mutated AML. The contradiction between prior experimental observation and our data in AML patients *in vivo* could be due to evolutionary biology of TP53 mutated AML in patients. Previous studies have shown that clonal hematopoiesis with TP53 mutation, without evidence of leukemia, is common in elderly individual ^29–33^. This suggests that in early stages of TP53 mutation, the mutated clone is actively cleared as it is formed. Hence it is possible that in the initial stages, TP53 mutation leads to activation of TE and immune pathways, enabling clearance of the mutated cells. But further clonal evolution *in vivo* via suppression of dsRNA may lead to escape from immune mediated attack of TP53 mutated cells, a hypothesis that needs to be investigated.

Recent report suggested that TP53 mutated AML are highly susceptible to treatment with hypomethylating agents ^34^, the mechanism of which is not known. In our study, despite low expression of TE and other non-coding RNA, the gene network analysis showed a moderately activated interferon gene pathway in TP53 mutated AML. Hypomethylating agents have been shown to activate the expression of TE and the downstream interferon leading to cancer cell death ^7,8^. We speculate that the specific gene expression signature of TP53 mutated AML (low dsRNA with active interferon gene expression) may make the leukemic cells vulnerable to further increase in dsRNA mediated activation of interferon and its resultant immune mediated cell death.

Our findings and previous work suggests that TP53 is a regulator of non-coding RNAs including TE. The mechanism of its regulation and whether it is a repressor or activator of TE expression *in vivo* in hematopoietic stem/progenitor cells and AML leukemic cells needs to be investigated.

Although methylation has been previously shown to regulate the expression of TE ^7,8^, our study showed very little association between mutations in methylation regulating genes (DNMT3A and TET2) and dysregulation of TE expression. However, we found strong association between mutations in TP53, NPM1, MLL and the cohesion complex. Mechanism of how these genes regulate TE expression will be needed.

In the past, assumptions have been made generalizing the function of TE as a whole without understanding their diversity. Most studies describe activated TE expression to be advantageous to the cancer cell by promoting genomic instability and hence cellular heterogeneity and cancer progression. However, recent studies describe their role in promoting cancer cell death via activating immune pathways ^7,8^. In this context, we recently showed that leukemic stem cells in AML, which are the most resilient to treatment, have suppressed TE expression ^21^. A recent study also reported that TE is suppressed in chemotherapy resistant cancer cells ^35^. Here, we similarly show that TE are suppressed in high-risk AML, particularly in TP53 mutated AML. These findings demonstrate that suppression of TE in cancer could possibly play a role in cancer evolution by protecting the cells from immune mediated cell death. TE constitutes a diverse group of transcripts, which likely have diverse function, as demonstrated by our network analysis correlating various TE biotypes with coding gene modules. It is likely that some of the TE that have active transpositioning activity promote tumorigenesis, whereas most others that have defective transpositioning activity perform other functions such as regulating coding gene expression and interferon activation via dsRNA recognition pathways.

This is the first study demonstrating the utility of TE expression signature in predicting prognosis in cancer. Our initial discovery using a large adult AML transcriptome database was validated in 2 independent cohorts, including a large pediatric AML cohort. We have identified 17 TEP transcripts that can be developed as a biomarker for prediction of survival in AML. Interestingly, the TE transcripts were associated with either good or bad prognosis and TE transcripts within the same class/type were associated with varying hazard estimates, indicating diversity within TE. For example, LTR34 and MER89, both from ERV1 biotype, were associated with good and bad prognosis, respectively. We predict that individual TE transcripts are as diverse as coding genes in their function.

Piggback derived 5 (PGBD5), which belongs to the DNA element family, was recently identified to be associated with poor prognosis in rhabdomyosarcoma ^6^. PGBD5 was then shown to exert its effect by interacting with HPRT1 and THAP9 genes. Similarly, studying the mechanisms of how the 17 TEP transcripts influences prognoses in AML may lead to novel understanding of the disease pathogenesis.

We propose an improved prognostic algorithm in AML utilizing mutational status along with expression of the 17 TEP. TP53 and NPM1 mutations by themselves conferred strong prognoses (poor and good respectively) and TEP expression information only marginally improved the risk classification. However, TEP significantly improved risk stratification of AML patients with mutations such as DNMT3A and FLT3 mutations. The utility of TEP in risk stratifying AML needs to be further validated using orthogonal assays in future studies.

Large adult and pediatric transcriptome data is available for multiple cancers in TCGA and TARGET. This study establishes the analytical foundation to investigate the role of TE in other cancers. Whether TEP identified in AML overlaps with other cancer and pattern of TE dysregulation observed with TP53 mutated AML is common across other cancers with TP53 mutation needs to be investigated.

## Supplement figure

**Supplement figure 1.**
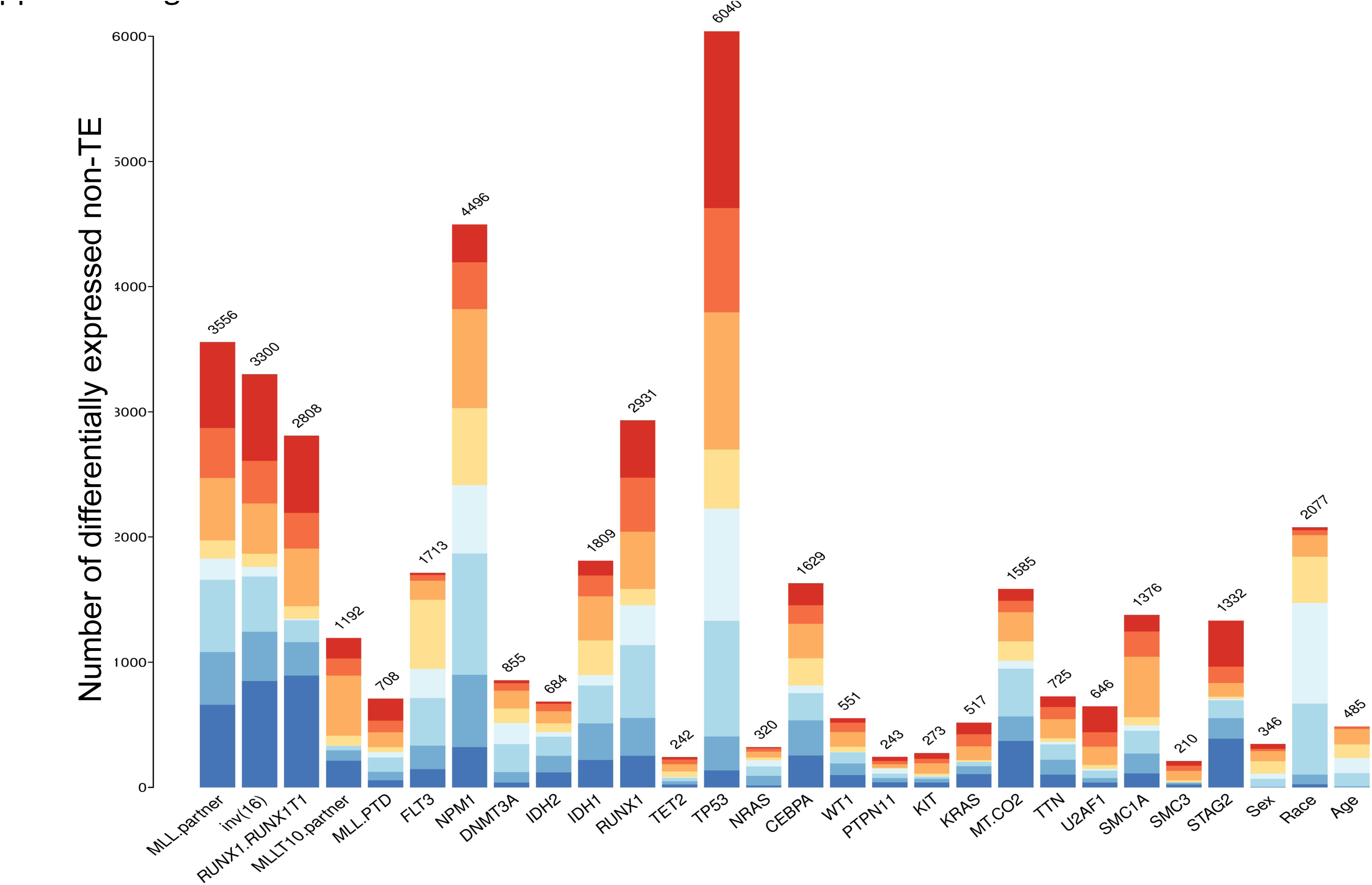
Similar to Figure 1, we examine the multiple regression measuring the non-TE ~ prevalent mutations. The prevalent mutations (5% prevalent) are on the x-axis. The y-axis is the total number of dysregulated non-TE for that corresponding mutational predictive factor used in the linear model. Only significant dysregulated non-TE were considered (hierarchical test, FDR method BH with significance level of 0.05).

**Supplement figure 2.**
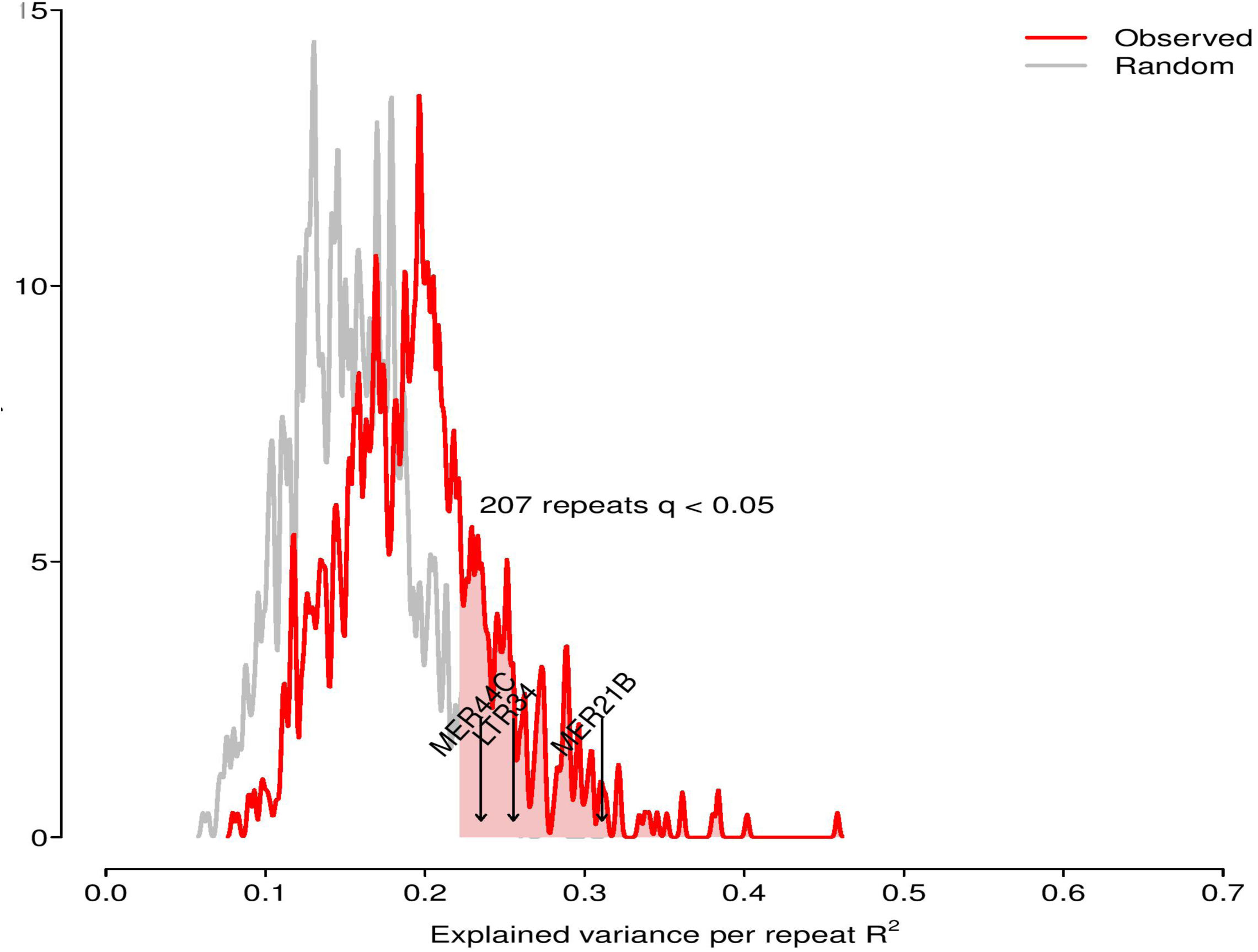
The x-axis is the explained variance per repeat elements using the true mutational profile as predictive features in a linear model (red). The y-axis is the density distribution range. The grey distribution is generated after randomly permuting the mutational profile. We've highlighted 3 TEP (table 1) in the critical region suggesting that some of the TE prognosticators are statistically altered in expression in this model.

**Supplement figure 3.**
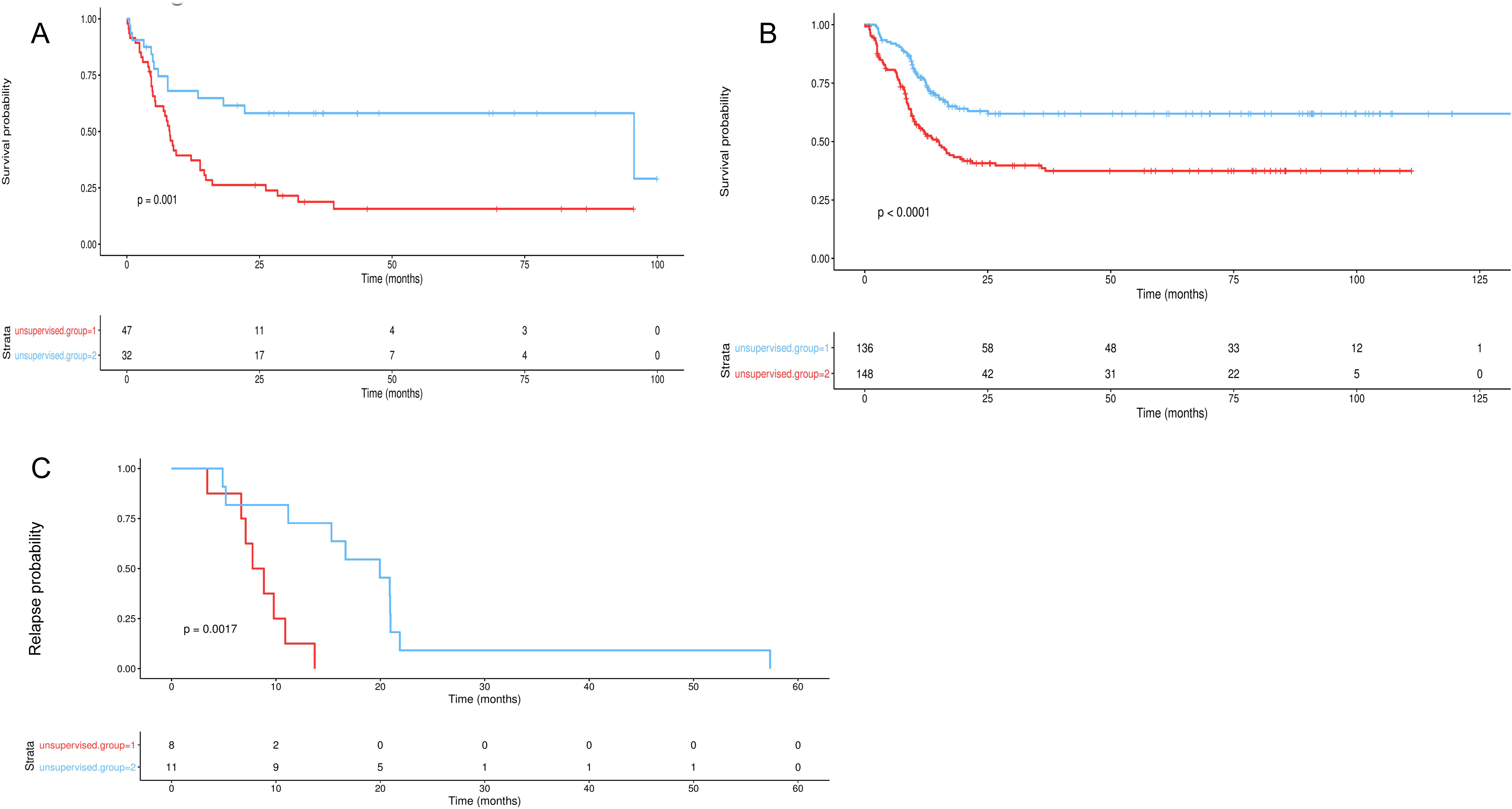
A) Survival analysis using 17 TEP in the TCGA test cohort (N=79). The y-axis is survival probability. The x-axis is Gme in months. Red is high risk classi?caGon, and blue is low risk. The groups were formed using Cox proporGonal hazards model (without applying any penalty in the model). PaGent unsupervised groups were formed using k-means clustering on the compound prognosGc summary values. B) Similar unpenalized model to A). Using Target (N=284), using 16 TEP. C) Similar unpenalized to A). ValidaGon in MSKCC (N=19) using 14 TEP. The y-axis is relapse probability.

**Supplement figure 4.**
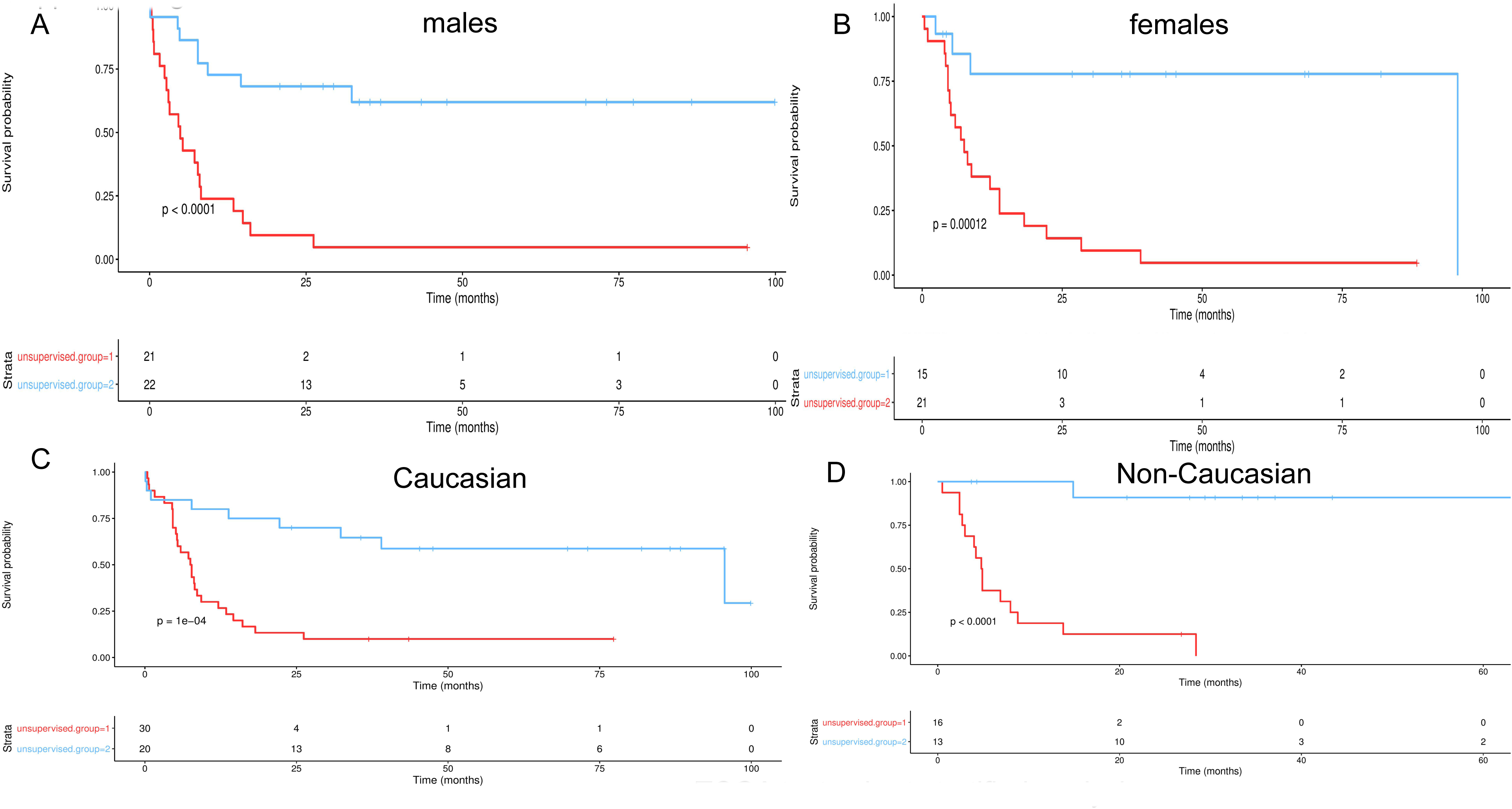
TCGA test cohort stratified analysis

**Supplement figure 5.**
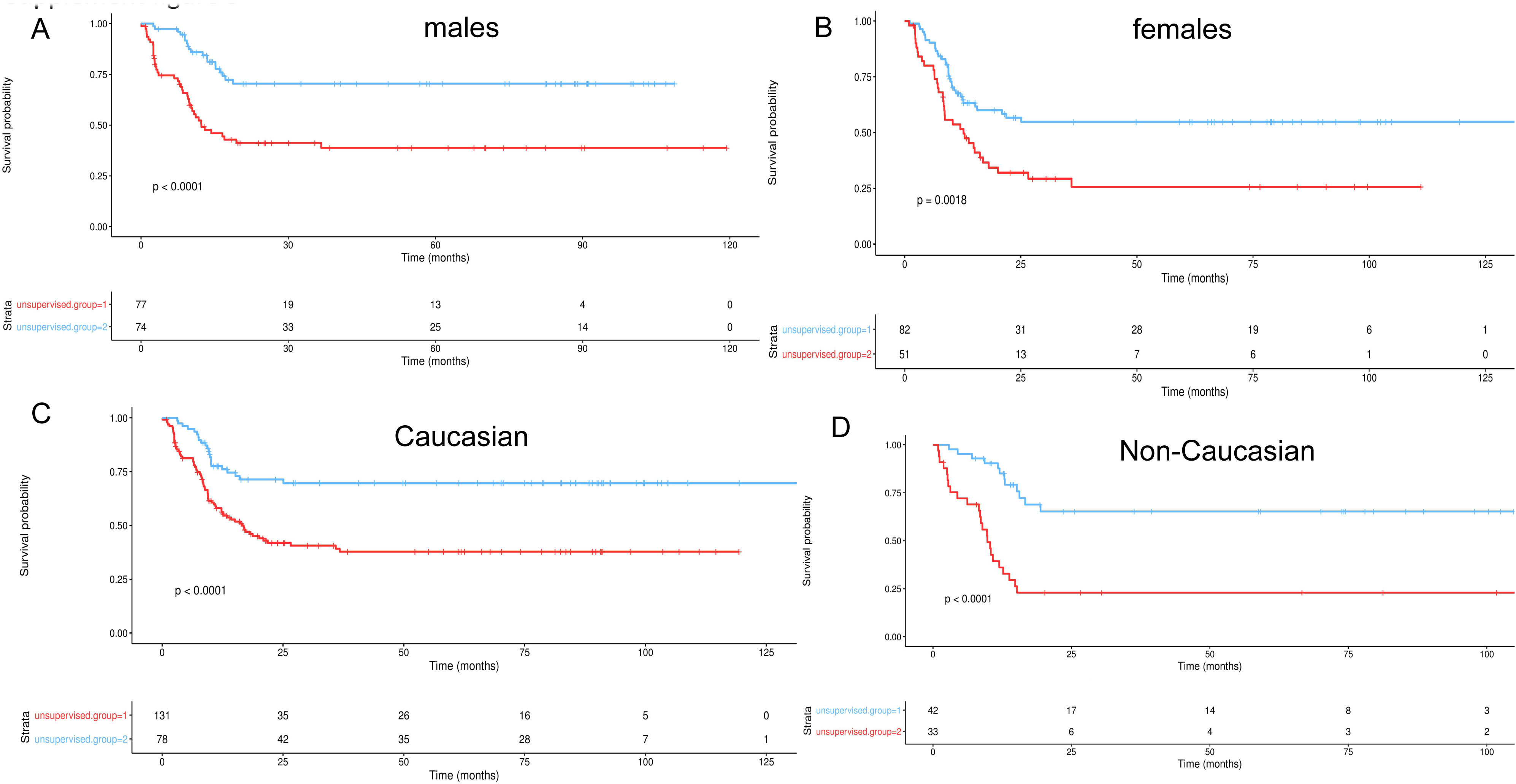
TARGET stratified analysis

**Supplement figure 6.**
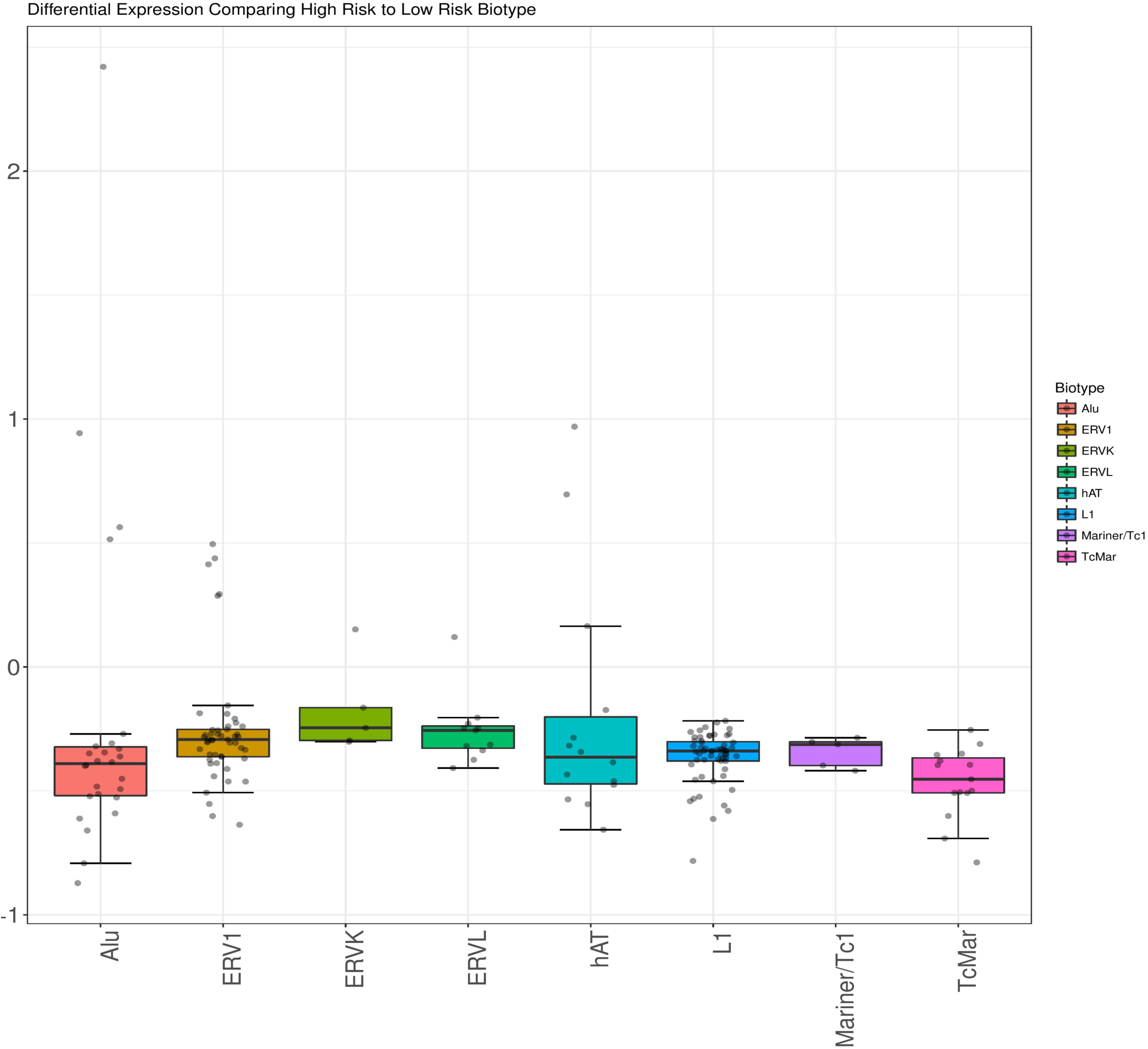
This compares TCGA full model (N=178) contrasting the differential expression using TEP classifications. The high risk patients are compared in a differential expression model to the low risk patients in a simple contrast (High.vs.Low). Differential expression was modeled using the high risk classification as the design matrix, and an empirical Bayes linear model was used to determine differential expression between TEs (adjusted p.value <0.05, adjust method is Benjamini-Hochberg). We observed that for all TE biotypes the low risk group has up-regulated expression in comparison to the high risk cohort. The box plot depicts the log-FC values on the y-axis calculated from the linear model after FDR filtering. The x-axis is the TE biotypes. We modeled all the TE transcripts simultaneously, and for each TE biotype calculated the summary statistics of the logFC computed from the model.

